# BindCompare: A Novel Integrated Protein-Nucleic Acid Binding Analysis Platform

**DOI:** 10.1101/2024.04.04.588140

**Authors:** Pranav Mahableshwarkar, Jasmine Shum, Mukulika Ray, Erica Larschan

## Abstract

**Summary:** Advanced genomic technologies have generated thousands of Protein-Nucleic acid binding datasets that have the potential to identify testable gene regulatory network (GRNs) models governed by combinatorial associations between factors. Transcription factors (TFs) and RNA binding proteins (RBPs) are nucleic-acid binding proteins regulating gene expression and are key drivers of GRN function. However, the combinatorial mechanisms by which the interactions between specific TFs and RBPs regulate gene expression remain largely unknown. To identify possible combinations of TFs and RBPs that may function together, developing a tool that compares and contrasts the interactions of multiple TFs and RBPs with nucleic acids to identify their common and unique targets is necessary. Therefore, we introduce BindCompare, a user-friendly tool that can be run locally to predict new combinatorial relationships between TFs and RBPs. BindCompare can analyze data from any organism with known annotated genome information and outputs files with detailed genomic locations and gene information for targets for downstream analysis. Overall, BindCompare is a new tool that identifies TFs and RBPs that co-bind to the same DNA and/or RNA loci, generating testable hypotheses about their combinatorial regulation of target genes.

**Availability and Implementation:** BindCompare is an open-source package that is available on the Python Packaging Index (PyPI, https://pypi.org/project/bindcompare/) with the source code available on GitHub (https://github.com/pranavmahabs/bindcompare). Complete documentation for the package can be found at both of these links.

## 1. Introduction

Spatio-temporal regulation of gene expression is a tightly regulated hierarchical cascade of interacting proteins binding to the genome and transcribing RNA molecules. Transcription factors (TFs) regulate RNA transcription from DNA, and RNA-binding proteins (RBPs) form the bulk of nucleic acid-binding proteins that control splicing and nuclear export. Multiple nucleic acid binding proteins function together to co-regulate specific sets of target genes, which are often called gene regulatory networks (GRNs)(Oksuz et al., 2023; Wang et al., 2022). High-throughput genomic technologies have generated hundreds of data sets that provide information regarding DNA and RNA targets of individual nucleic acid binding proteins, helping decipher the combinatorial function of TFs and RBPs in regulating specific GRNs (Salma et al. 2023; Martin et al. 2023; König et al. 2012). However, user-friendly analysis tools cannot generate high-throughput comparisons between multiple protein-nucleic acid binding datasets to determine common and unique targets.

We present a novel command line and interactive tool for analyzing coordinated protein binding to DNA and RNA called BindCompare that: i) compares multiple protein-nucleic acids binding datasets to identify interacting pairs of factors in an unbiased way; ii) identifies common targets between selected protein pairs identified by the user. Thus, BindCompare will enable the identification of inter-dependent and combinatorial RNA/DNA binding, generating new hypotheses about how single and dual RNA and DNA binding factors impact gene regulation. To ensure user accessibility, BindCompare has a user-friendly graphical user interface (GUI) that can be used to study any organism.

## 2. Method

The main inputs to BindCompare are peak-called Browser Extensible Data (BED) files (**Figure 1A**). The first module in BindCompare, **bindexplore**, identifies combinatorial binding patterns across the genome (**Figure 1B**) to be further studied with **bindcompare** (**Figure 1C**). **bindcompare** profiles all overlapping nucleic acid binding regions for any pair of two proteins and extracts related sequence and gene information. These annotated results allow downstream analysis through BindCompare’s **comparexp** module and other tools (**Figure 1D, E**).

**Figure 1.**
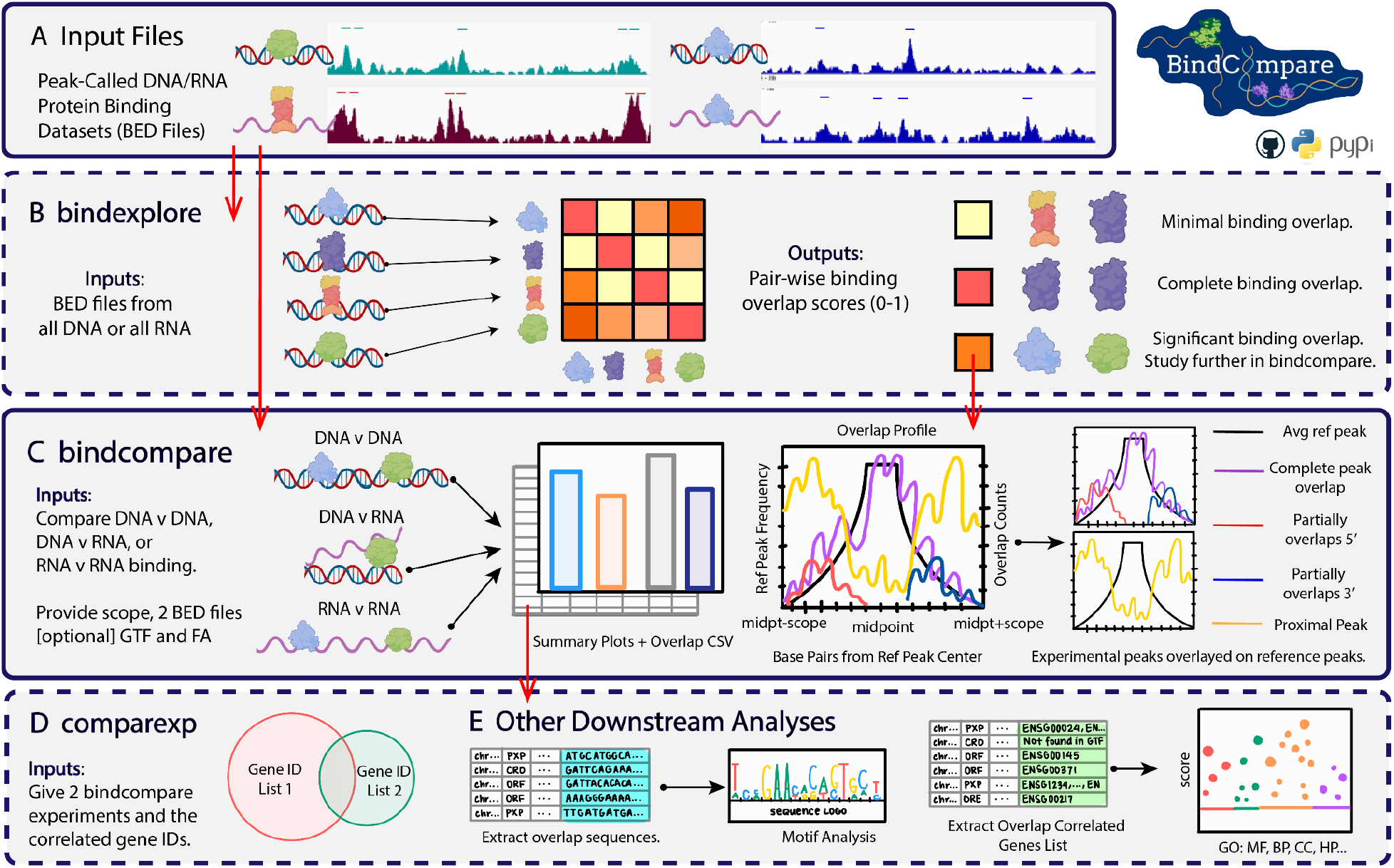
Schematic of the BindCompare workflow. **A**. Inputs to BindCompare are peak-called binding datasets from experiments such as ChIP-seq (DNA binding) and eCLIP (RNA binding) in BED format. Bars above high-intensity signals correspond to hypothetically called peaks. **B**. If working with protein binding (BED files) to only DNA or RNA, the bindexplore function can identify pair-wise binding interactions between N data sets, helping identify candidate proteins to study with the **bindcompare** function. **C. bindcompare**, the core function, performs comprehensive binding analysis between two sets of BED files (DNA/DNA, RNA/RNA, or RNA/DNA), producing a diverse array of binding profiles and other outputs. **D. comparexp**, when paired with the gene GTF file, allows users to identify factors with overlapping binding sites. **E**. Gene lists and sequences correlated with binding overlaps can be used for Gene Ontology and motif analysis.

Protein-nucleic acid assays, including eCLIP, ChIP-seq, and CUT&RUN (Van Nostrand et al. 2016; Skene and Henikoff 2017) yield peak-called BED files listing genomic binding locations. Platforms such as Model-based Analysis of ChIP-Seq (MACS2) or PureCLIP (Gaspar 2018; Krakau, Richard, and Marsico 2017) are also used for peak calling.

### 2.1 bindexplore - Multifactorial Binding Analysis

Users can input ‘N = total number’ of peak-called BED files for DNA or RNA into the **bindexplore** module, producing a heatmap of co-binding activity across all pairs of proteins (**Figure 1B**). The method first splits the genome into user-defined bins of default size 1000 bp (**Table 1, Figure S1;** see **supp. file 1, section 1.2** for further details). All binding sites are then mapped into the bins. For all N^2^ protein pairs, pairwise intersections of binding-correlated bins are determined by choosing one protein as the reference and overlaying the other. The correlation value for each pair equals the number of times both proteins bind in the same bin divided by the number of times the reference protein binds, providing a normalization (**supp. file 1, Equation 1**).

**Table 1.**
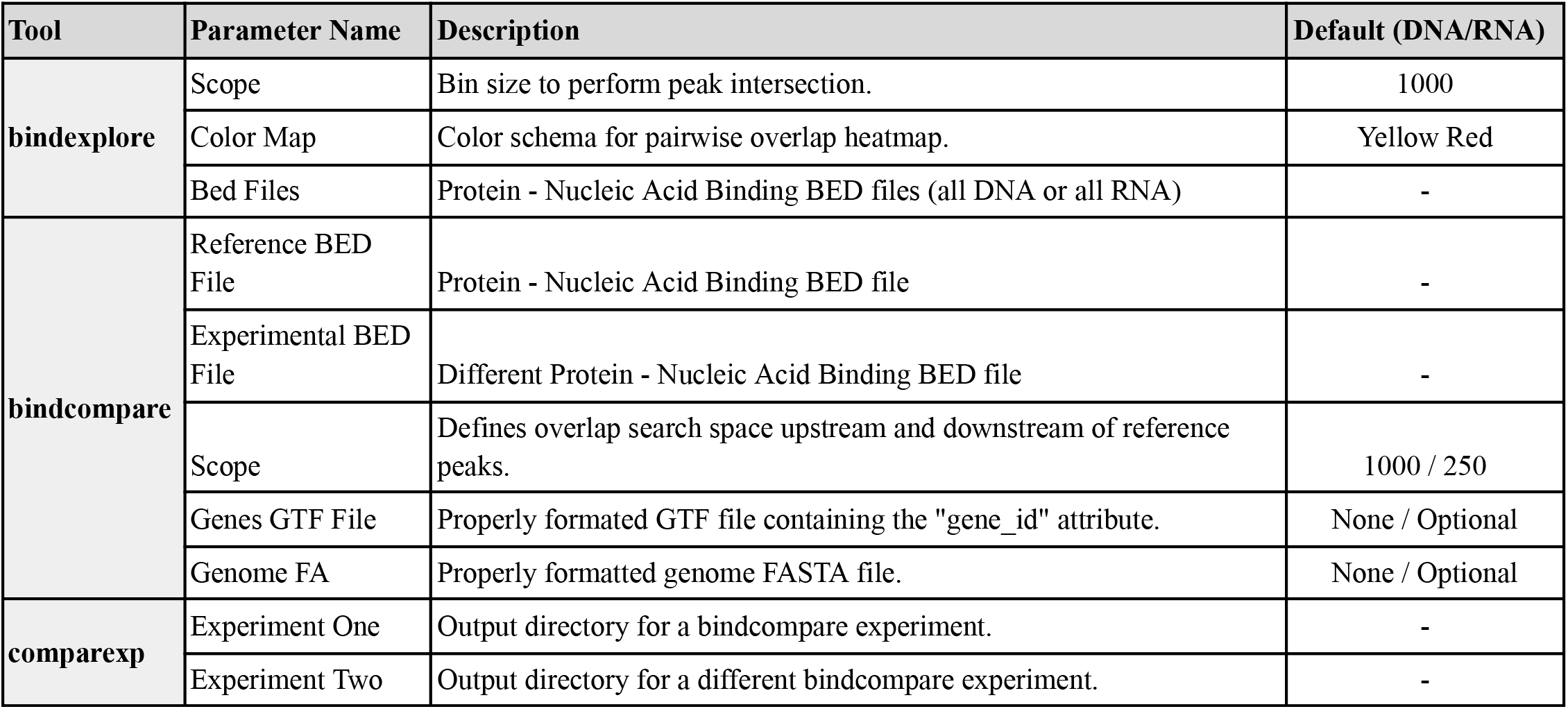
Parameters for BindCompare modules.

| Tool | Parameter Name | Description | Default (DNA/RNA) |
| --- | --- | --- | --- |
| **bindexplore** | Scope | Bin size to perform peak intersection. | 1000 |
|  | Color Map | Color schema for pairwise overlap heatmap. | Yellow Red |
|  | Bed Files | Protein - Nucleic Acid Binding BED files (all DNA or all RNA) | - |
| **bindcompare** | Reference BED File | Protein - Nucleic Acid Binding BED file | - |
|  | Experimental BED File | Different Protein - Nucleic Acid Binding BED file | - |
|  | Scope | Defines overlap search space upstream and downstream of reference peaks. | 1000 / 250 |
|  | Genes GTF File | Properly formatted GTF file containing the "gene_id" attribute. | None / Optional |
|  | Genome FA | Properly formatted genome FASTA file. | None / Optional |
| **comparexp** | Experiment One | Output directory for a bindcompare experiment. | - |
|  | Experiment Two | Output directory for a different bindcompare experiment. | - |

The overlap between a pair of proteins targeting common nucleic acid target sequences is represented by a correlation value between 0-1 (yellow-red) (**Figure 1B, Supp. File 2, Figure S5A, B**). This process reveals candidate co-regulators that can be further analyzed with **bindcompare** (**section 2.2, supp. file 1, section 1.3**).

### 2.2 bindcompare - Pairwise Binding Analysis

With the **bindcompare** module, two sets of protein-nucleic data chosen by the user can be compared, including the following possible comparisons: i) protein-DNA and protein-DNA, ii) protein-RNA and protein-RNA, or iii) protein-DNA and protein-RNA **(Figure 1C, Table 1**). In addition to these two inputs, the script requires a scope, the range at which the binding sites should be called as overlapping, and optional genome fasta and genes GTF files. The scope accounts for the co-regulatory interactions that sometimes occur at neighboring genomic locations due to the resolution of the experimental technique and potential nucleic acid folding. As the scope is increased, more overlaps are detected, though at a decreasing rate. The default values for **bindcompare** were chosen to reflect the inflection points where the rate of new overlaps detected stabilizes as the scope increases (**Figure S2, Table 1;** see **supp. file 1, section 1.4**).

**Bindcompare** uses the provided scope to look for overlapping binding sites upstream and downstream of each reference peak. These overlaps are then classified into four categories: 1) complete reference overlap (CRO); 2) Overlap Reference End or 3’ overlaps (ORE); 3) Overlap Reference Front or 5’ overlaps (ORF); 4) proximal peak (PXP) (**Supp. Methods 1.3**). This process is optimized using interval trees to store binding loci (Ang and Tan 1995; Halbert and Tretyakov 2020). Then, the script outputs a visualization of the binding profiles of these proteins, plotting the average reference peak and the frequency of overlapping experimental peaks across the scoped region. In addition, the script provides the distribution of overlap types and visualization of binding profiles specific to each chromosome. The binding profile contains four colored lines over the average reference peak, each correlating to one of the four categories of overlap described above. The y-axis indicates the number of times that this specific overlap event occurred at a position over the scoped region. Additional plots produced by **bindcompare** place co-regulatory activity in the context of overall binding patterns for both protein-binding datasets (**Figure S7, Table S1c-g**). Further, **bindcompare** can directly compare DNA and RNA binding for specific TFs (**Figure 1C**), which is poorly understood despite most TFs binding to DNA and RNA (Oksuz et al. 2023).

### 2.3 comparexp and Downstream Analyses

The final module in BindCompare, **comparexp**, enables users to determine the impacts of genetic, RNAi, or small-molecule perturbations on locations of protein-nucleic acid binding (**Figure 1D, supp. file 1, section 1.5, Table 1**). **comparexp** identifies unique protein-nucleic acid interactions occurring in only one experimental condition and overlapping targets that occur in multiple conditions.

Furthermore, given a genome file or genes GTF file, **bindcompare** outputs all genes associated with the bound sequences that change after a perturbation. These outputs can be input into popular Gene Ontology tools, including GProfiler2 (Kolberg et al. 2020) and ShinyGO (Ge, Jung, and Yao 2020). Finally, sequence information can be studied using the MEMESuite (Bailey et al. 2015) (**Figure 1E**).

### 2.4 Using BindCompare and Comparing to Existing Methods

BindCompare can be used either through i) the command line or ii) an interactive GUI. Users can install the tool using the Python Package Index (https://pypi.org/project/bindcompare) or GitHub (https://github.com/pranavmahabs/bindcompare). To increase the accessibility of the BindCompare pipeline to users who do not use the command line, the scripts can be run through a Python CustomTkinter GUI interface ((Schimansky 2024); **supp. file 1, section 1.6**). The GUI platform and extensive documentation help to analyze the visual outputs. A detailed user interpretation guide can be found on the home documentation page.

Bedtools is a multi-functional bioinformatics tool that enables users to apply set theory functionality on BED files and is broadly used in bioinformatics analysis pipelines (Quinlan and Hall 2010). When comparing the overlap events detected by **bedtools intersect** and **bindcompare, bindcompare** reports 146.08% additional events in comparison to **bedtools intersect** (**Figure S4**). This difference can be attributed to **bindcompare** searching for overlap events in the scoped region bounding each reference peak and not only within the binding region as **bedtools intersect** does. This ability to quickly profile upstream and downstream regions is unique to BindCompare. It provides a useful output for biologists because biologically relevant overlaps between binding events for different factors rarely align precisely with each other.

Overall, BindCompare is different than existing tools for five key reasons: 1) BindCompare allows users to rapidly generate new hypotheses by comparing a large number of data sets to each other; 2) BindCompare allows users to compare DNA and RNA binding profiles of the same protein to each other; 3) BindCompare broadens the definition of a peak overlap to profile proximal regions; 4) BindCompare does not require users to have a background in scripting due to an intuitive GUI interface; 5) finally, BindCompare can be installed easily for any architecture (including Apple Silicon), a significant barrier for current bioinformatics packages (see **supp. file 1, section 1.8** for details).

## 3. Results

### 3.1 Identifying common targets of multiple proteins that bind to nucleic acids

Using the **bindexplore** module (**Figure 1B**) of BindCompare, we first show how the user can identify common nucleic acid targets shared by multiple proteins. Protein-DNA binding ChIP datasets and protein-RNA binding eCLIP datasets for 16 human RBPs from ENCODE project phase III (GSE77634) were separately run through **bindexplore**, generating two correlation matrices (**supp. file 2, Figure S5A, B; supp. file 3, Table S1a, b**) for pairwise comparison between all sixteen RBPs’ DNA and RNA binding. We found that RBP HnRNPA1 binding to RNA has a high correlation matrix score (**CMS**) (>0.50) with the following reference RBPs binding to RNA: Sfpq (0.57), Fus (0.63), Taf15 (0.64), and Hnrnpul1 (0.69), indicating that they share many common targets. In the literature, all five RBPs are known to be associated with neurodegenerative diseases like ALS (Amyeloid Lateral Sclerosis) and regulate RNA splicing (Gordon et al. 2021; Xue et al. 2020). When we overlapped Fus, Sfpq, Hnrnpul1, and Taf15 binding on RNA with HnRNPA1-RNA as a reference, CMSs were <0.50, with reciprocally high CMSs between each other. This suggests that the first four proteins also have HnRNPA1-independent common targets. Thus, **bindexplore** generates hypotheses about the combinatorial function of factors by defining the fraction of common targets between specific protein pairs, which could be further tested using other experimental approaches. The **bindexplore** correlation matrix for the DNA binding (ChIP) for the same set of RBPs (**supp. file 2, Figure S5B; supp. file 3, Table S1b**) shows that Taf15 has more common DNA targets with Ptbp1, Srsf1, and U2af1 despite having a low score for common RNA targets (**supp. file 2, Figure S5A, B**). Therefore, **bindexplore** can be used to build a network of combinations of TFs and RBPs targeting specific chromatin loci at the DNA and/or RNA levels to define a putative regulatory network at specific genomic locations containing target genes of interest.

Next, the user can run the **bindcompare** module (**Figure 1C**) of the BindCompare tool to make pairwise comparisons of specific protein pairs of interest using high correlation matrix scores as a guide. Here, we compared HnRNPA1 pairwise to RBPs with high correlation matrix scores defined by **bindexplore** and identified common targets shared by HnRNPA1, Sfpq, Fus, and Taf15 (**supp. file 3, Table S1c**). **bindcompare** also compares two different peak distributions (**supp. file 2, Figure S6A-D**) by defining the following categories of overlapped peaks: 1) complete overlap between reference (Rbfox2-DNA) and overlapped (Ptbp1-DNA) protein-DNA binding regions (**CRO)**; 2) 5’ front partial overlap with reference (**ORF**); 3) 3’ end partial overlap with reference (**ORE)**; 4) proximal peaks (+ 1000bp scope) to reference. Thus, the user can identify genome-wide shared targets for each pair of factors.

### 3.2 Identifying relative binding of protein pairs to DNA and RNA: regulation of co-transcriptional RNA processing

We show that **bindcompare** identifies the extent of co-binding to nucleic acids between different proteins and generates testable hypotheses about the co-transcriptional regulation of RNA processing (Bentley 2014; Cordiner et al. 2023). The Taf15 and Fus proteins regulate transcription and RNA processing (Schwartz, Cech, and Parker 2015), but their shared DNA and RNA targets remain unknown. We used **bindcompare** to show that Taf15 and Fus share common DNA and RNA targets (**supp. file 2, Figure S7A-B**) and that one-fourth of their DNA targets are also associated with RNA (**Supp. File 2, Figure S7C-D**). Therefore, these proteins may regulate co-transcriptional RNA processing at a specific subset of their binding sites, which **bindcompare** identified (**supp. file 3, Table S1d-g**).

Furthermore, **bindcompare** identifies the relative positions where RBPs/TFs bind to common nucleic acid targets. The frequency plot (overlap profiles) (**Figure S7E-H)** outputs of **bindcompare** show that Taf15 binds to RNA within 100 nucleotides of Fus binding at shared RNA targets. In contrast, DNA targets bind side-by-side within a scope of 250 bp, as shown by the higher frequency of partially overlapping peaks at both ends of the reference peak (**Figure S7E-F)**, with both Taf15 and Fus connecting RNA to DNA within 250 bp (**Figure S7G-H)**.

The user can also do individual chromosome-specific binding analysis with BindCompare. We compared the binding of the male X-chromosome enriched *Drosophila* transcription factor CLAMP at genomic locations on DNA (# GSE220053) with its RNA targets (# GSE205987) in male cells (Ilik et al. 2013; Soruco et al. 2013; Ray et al. 2024). Using BindCompare for individual chromosome-wise overlap profiles, we show that CLAMP-DNA peaks overlapping with RNA peaks are more frequently distributed on X-chromosomes than autosomes (**supp. file 2, Figure S8**), consistent with literature demonstrating that CLAMP is enriched on the male X-chromosome and interacts with RNA on the male-X.

### 3.3 Identifying unique/common protein-nucleic acid binding targets between different experimental conditions

Many studies focus on the dependence of the binding of a single RBP or TF to its targets on another RBP/TF. Using **comparexp (Figure 1D)**, we compared common DNA targets for the interacting partner’s *Drosophila* TF CLAMP and the *Drosophila* RBP RNA helicase MLE (Maleless) (Tikhonova et al. 2024) in the presence and absence of TF CLAMP (**supp. file 2, Figure S9**). **comparexp** determined that at 537 CLAMP binding loci, MLE targets DNA only in the absence of CLAMP, i.e., CLAMP inhibits MLE binding to DNA at these ectopic binding loci. Thus, **comparexp** determined how the chromatin binding of one RBP, MLE, is promoted or inhibited at different loci by its binding partner, the TF CLAMP.

Overall, we show that BindCompare is a new tool that provides essential information for identifying and studying the combinatorial binding of TFs and RBPs. Moreover, BindCompare has the potential to reveal novel mechanisms regulating co-transcriptional RNA processing because it is the only available tool that simultaneously measures DNA and RNA binding to chromatin.

## Author Contributions

PM and MR conceived the project idea. PM implemented the BindCompare platform under the guidance of MR and EL. JS and PM ran analyses on the BindCompare platform and developed figures. PM and MR wrote the manuscript, and EL provided revisions.

## Data Availability

BindCompare is an open-source package that is available on the Python Packaging Index (PyPI, https://pypi.org/project/bindcompare/) with the source code available on GitHub (https://github.com/pranavmahabs/bindcompare). Complete documentation for the package can be found at both of these links.

