## Supplemental File 1: Methods for "BindCompare: A Novel Integrated Protein-Nucleic Acid Binding Analysis Platform"

### Supplementary File 1

#### 1.1: Data

**BED Files and Input Data:** The input to BindCompare are peak-called Browser Extensible Data (BED) files. These can be obtained from experiments including, but not limited to, enhanced crosslinking and immunoprecipitation (eCLIP), chromatin immunoprecipitation sequencing (ChIP-seq), and cleavage under targets & release using nuclease (CUT&RUN). The raw sequence files can be processed using documented pipelines and peak-called using programs such as Model-based Analysis of ChIP-Seq (MACS) or PureCLIP. This will produce peak files listing a peak-called binding site for the protein of interest on each row, listing the chromosome, the start, and the stop positions.

Frequently, these experiments are conducted using multiple replicates, as is the case with the hg19 eCLIP data presented here. Consensus peaks can be resolved in preprocessing or by selecting only peaks that appear at some minimum threshold across the replicates. This can be done utilizing Bedtools and common command-line programs. Here, eCLIP data from GSE77634 contained two replicates, and consensus peaks were chosen by selecting sites found in both peak-called replicate files.

#### 1.2: bindexplore

The input to **bindexplore** is N peak-called BED files that list peaks belonging to the same nucleic acid (i.e., all DNA binding peaks). The **bindexplore** script takes in a scope, or bin size, and splits the genome into bins of size scope. This user-defined scope can be larger or smaller depending on the desired distance to be captured between co-localizing proteins. Then, **bindexplore** uses a BedProcessor object to iterate through the binding sites and place each binding site into its corresponding bin by dividing the start site by the scope size. Unlocalized contigs, scaffolds, and improperly labeled rows should be discarded when processing binding sites. The bin IDs for binding sites for each provided file are then stored in lists split by chromosome. All BedProcessor objects are managed by a ProcessManager object that takes in a list of BED files.

Once all files have been processed, the ProcessManager performs pairwise intersections to create a correlation matrix *M*. For a matrix value $M_{i,j}$, the binding site dictionary is retrieved for BED files *i* and *j*. Then, the correlation matrix is built using Equation 1. These calculations allow for redundancy/repeated appearance of bins. This ensures that bins that contain multiple peaks are properly accounted for when calculating correlation scores.

$$M_{ij} = \frac{\sum_{\text{chr}=a} \text{Number of Binding Sites in } j \text{ found in } i \text{ or on chr } a}{\text{Number of Binding Sites in } j}$$

**Equation 1 bindexplore correlation matrix M calculation**: The sum over chromosomes is taken for the number of binding sites in BED j found in BED i. This is done using the NumPy isin() function. Then, the sum is divided by the number of binding sites in BED j to normalize the values.

Every individual BED file will have variable numbers of peaks because some factors bind less often with high specificity and others very often with low specificity. If we were to present the results of **bindexplore** using raw counts (not normalizing with the division term in Equation 1), then the intersection scores would skew towards the nucleic-acid binding proteins with low specificity. Scaling all counts by the number of reference peaks shows what proportion of a protein's peaks overlap with another set of peaks. This makes the correlation score more informative and allows easier comparisons across protein pairs.

Then, these values are plotted using Matplotlib's show () function, treating the correlation matrix as an image. A color bar shows the range of values from 0 to 1. The plot and raw values are saved in CSV format.

In **Figure S1**, we examine the relationship between scope and the interpretability of **bindexplore** results. The 30 bp scope is highly similar to a direct intersection of BED file peaks and yields broadly lower values. Intermediate scope choices surface more complex relationships. At a higher scope of 5000bp, the heatmap oversaturates.

**1.3: bindcompare**

**Core Functionality and Backend:** There are seven inputs to **bindcompare**, two of which are optional. A reference BED file, an experimental BED file, a scope, and the experiment name/output directory are required. Optionally, users can provide a genome fasta file and a gene GTF file that follows standard formatting. Once all the inputs are received, two Bed objects are created that load and process all binding sites in each BED file, storing the information by chromosome. Unlocalized contigs, scaffolds, and improperly labeled rows are disregarded in chromosome processing. Additionally, the reference peaks are expanded to range from *[midpoint - scope, midpoint + scope],* making each reference peak the same size while retaining the original peak start and stop in metadata.

For each chromosome, the binding information is stored utilizing interval trees. This allows for a logarithmic time complexity in retrieval. Then, a BindCompare object is created, taking in the reference and experimental data, allowing for the overlap through the compare_binds function. Within, the compare_chrom_bind_it() function is called for each chromosome. This function iterates over each experimental binding site and utilizes the interval tree for efficient retrieval (O(log(N)) of all reference binding peaks that overlap this experimental peak. Then, each of these overlaps is categorized into four categories: complete reference overlap (CRO), overlap reference end (ORE), overlap reference front (ORF), and proximal peak (PXP). These overlaps are defined in Equation 2 for a reference peak *chrA-b:c* and experimental peak *chrD-e:f*.

$$\begin{aligned}
\text{CRO} &= \mathbb{1}_{A=D}\left((e \geq b \wedge f \leq c) \vee (e \leq b \wedge f \geq c)\right) \\
\text{ORE} &= \mathbb{1}_{A=D}\left((b \leq e \leq c) \wedge f > c\right) \\
\text{ORF} &= \mathbb{1}_{A=D}\left(e < b \wedge (b \leq f \leq c)\right) \\
\text{PXP} &= \mathbb{1}_{A=D}\left(\left(\frac{b+c}{2} - \text{scope} < f < b\right) \vee \left(c < e < \frac{b+c}{2} + \text{scope}\right)\right)
\end{aligned}$$

**Equation 2 Mathematical definitions of overlap categories**: A CRO entails the experimental peak being completely contained by the reference peak. ORE and ORF entail the experimental peak partially overlapping on either side of the reference peak. Finally, PXP entails the experimental peak overlapping with the extended scoped region but not the original reference peak itself. For all, the chromosomes must match.

Every overlap found is added to a running set of all overlaps found thus far. The scoped region can be represented as *[-scope, scope]*. The region over this domain for which the overlap occurs is also stored. For example, if the overlap occurred from chrX:900-1400 for a peak chrX:1100-1500 and a scope of 1000, the positions *[-400, 100]* would be added for this overlap as these are the positions over the reference region that the overlap occurred. These values will be used later in developing the overlap profiles.

In this process, relevant metadata, including the number of unique overlaps, total overlaps, and proximal peaks, is counted for each chromosome. Then, all of this information is aggregated by chromosome in a **bindcompare** dictionary that will be used for downstream visualization.

**Overlap Profiles:** The overlap profile is the main visualization produced from **bindcompare**. This visualization from the aforementioned **bindcompare** dictionary retrieves the overlap frequencies or counts over the reference domain of *[-scope, scope]*. At each position, the number of overlaps of each type is counted. For example, at position *x,* a count of 30 for CRO entails that 30 experimental peaks yielded a complete reference overlap at position *x*. Then, these values are plotted as a line over the x-axis domain of the domain of *[-scope, scope]*. The purple line is CRO, blue for ORE, red for ORF, and yellow for PXP. The right-hand side y-axis provides the counts for the number of experimental overlaps of that type at each position across the scoped reference domains. In black, the average peak of the reference peaks is plotted with the frequency of reference peaks occurring at each position over the scoped region. This plot tends to be around 1 towards the central region of the domain.

Because all the required computations were performed at the chromosomal level, another plot showing the overlap profiles at the chromosomal level is also generated. The full version of the plot is created using Matplotlib's line plots. The second version utilizes the same line plots using subplots to place all chromosomes' overlap profiles in the same image.

**Additional Outputs:** Utilizing the metadata created from the overlap of the binding data, several additional plots can be created. First, two plots show the distribution of different overlap types in a stacked barplot (barh in Matplotlib) and a pie chart. Then, another bar graph is created that provides the number of experimental binding peaks, the number of unique overlaps (CRO, ORE, ORF), the number of overlaps, the number of proximal peaks (PXP), the unique number of proximal peaks, and the unique reference peaks identified in overlapping binding events. An event CSV is generated that lists every single overlap event documented. However, because a reference peak can overlap with multiple experimental peaks, these rows are collapsed into one row to avoid repeating information. Then, if a genes GTF file is provided, the correlated genes are identified in these regions in addition to the sequence of the reference peak if the genome FA is provided. The sequence information is also stored as a sequence.fa file. This information is also stored separately in four CSVs that are split based on overlap type. Finally, a summary file repeats some of the above metadata and prints a space-separated list of identified genes.

The genome fasta and gene GTF files are provided for users through BindCompare installation.

#### 1.4: Benchmarking bindcompare

To quantify the robustness of bindcompare to the choice of scope, **bindcompare** was run on the four sets of data 1. Fus DNA vs Fus RNA binding, 2. Taf DNA vs Taf, 3. Fus DNA vs Taf DNA binding, and 4. Fus RNA vs Taf RNA binding using a broad range of 26 scopes (**Figure S2**). Across all four comparisons, the runtime remains fairly consistent across all scope choices with minor fluctuations. As scope increases, there is a subtle increase in runtime, though never exceeding 5 seconds from the minimum runtime (8.69% of total runtime on average). Instead, fluctuations in bindcompare runtime can be attributed to the number of reference peaks and overlap events detected. Interestingly, beyond scope size 1000bp for DNA overlaps and scope size 300bp for RNA overlaps, the number of new overlapping events detected plateaus, suggesting potentially significant overlaps appear within these ranges. Therefore, we suggest 1000bp as the default scope for most comparisons and a scope of 250bp for RNA binding experiments.

All peaks and gene GTF information are stored in interval trees - an optimized data structure that allows for logarithmic time data accession when searching for overlaps. This ensures performant runtime in dataset intersection. Regardless, a larger dataset (i.e., more peaks or a larger genome) will take more time to query. This result can be seen in the experiments performed; when the

RNA bindcompare for RBPs FUS and TAF15 RNA is benchmarked in Figure S2C, the runtime is around 10 seconds longer than the runtime for the DNA comparisons in Figures S2A, S2B, and S2D. In particular, the DNA datasets had a combined 28781 peaks while the compared RNA datasets had around 500% more at 176281 peaks. This significant difference in dataset size only corresponds to an average 18.56% increase in runtime (which can be attributed to the interval tree-based data storage).

Altogether, these results show that bindcompare maintains fairly robust runtimes when the scope is changed in extreme values and that differences in runtime can be attributed to the actual data being tested.

**1.5: comparexp**

Given two **bindcompare** output directories that utilized a genes GTF file, this script opens the summary file and pulls the correlated gene lists for each **bindcompare** experiment. Then, a weighted Venn diagram (venn2 in matplotlib_venn) is created to show the intersection of these gene lists. The actual genes in each category are also printed in a summary file.

**1.6: BindCompare Visual Interface**

Accessibility is key to the usability of the BindCompare suite. Run bindlaunch in the command line to use the interface, and the visual interface will be launched. This is done through a CustomTkinter app built on top of Tkinter through the CustomTkinter package. On the left-hand side of the app, the user can be redirected to the home page for BindCompare, change the font size, and change the appearance mode. Additionally, the user can launch a sub-window to run **comparexp**. A rendering of the BindCompare GUI can be seen in **Figure S3**.

**Using BindCompare through a Python GUI:** In the central portion, the user can run **bindcompare** by selecting the files they would like to use for each input parameter through the file prompt. This decreases the user's required knowledge of file paths through the command line. Then, by clicking run, **bindcompare** is run as a subprocess in the code's backend. This changes the status bar in the GUI to indicate that **bindcompare** is running. Once complete, the status bar will update again, and any output produced by the script will populate within the GUI. These outputs could include potential error messages (i.e., invalid file input). Finally, once **bindcompare** has been run on the right-hand side, the user can load and visualize a compressed version of each plot. The user can find relevant information in the help text section that interprets different visualizations and plots. This complements a user interpretation guide that exists in the GitHub documentation.

In the **comparexp** window, the user selects two **bindcompare** directories. Once they click run, the weighted Venn diagram appears. The summary file is also loaded into a text box below, containing the space-separated gene lists in each of the three categories within the Venn diagram.

**Other Downstream Analyses:** If a gene GTF is provided, **bindcompare** and **comparexp** will produce space-separated gene lists. These can be fed into gene ontology programs such as ShinyGO or GProfiler2. Further, motif analysis can be run through FIMO, STREME, or other MEME Suite programs using the sequences.fa file produced by **bindcompare** if a genome fasta file is provided.

We used ID converters found on BioTools.fr to convert Fbgn (Flybase Gene IDs) and ENSEMBL Gene IDs to Gene Symbols before downstream analysis.

**1.7: Comparing BindCompare to Existing Methods**

Only two other tools, **Bedtools,** and **DiffBind,** occupy a space similar to BindCompare. Bedtools is a multi-functional bioinformatics tool that enables users to apply set theory functionality (such as merges, intersections, count, etc.) on BED files. As a result, the Bedtools platform is broadly used in bioinformatics analysis pipelines, including Dedtools' merge and overlap functions. Diffbind is a popular Bioconductor (R) package designed for thorough differential binding analysis across conditions or treatments within ChIP-seq and ATAC-seq experiments. Statistical tools and visualizations in the package aim to reveal specific sites of differential binding and their functional associations.

BindCompare has several unique features compared to these platforms. First, our platform is designed to comprehensively analyze all binding interactions, introducing the concept of an overlap profile. This unique visualization shows nucleotide position-based visualizations that compare binding positions at the genome and chromosome levels. Combined with the combinatorial analysis possible with bindexplore, BindCompare allows the user to develop testable hypotheses for co-regulatory binding functions. Furthermore, BindCompare can specifically compare protein-DNA with protein-RNA interactions, which is an important functionality because most transcriptional regulators have recently been shown to bind both DNA and RNA.

The existing **bedtools intersect** function performs the defined task of directly intersecting two sets of peaks. However, co-regulatory functions and biological behavior can also be captured by examining the binding activity in the immediate neighborhood of a binding site. BindCompare, by utilizing the scope, allows users to visualize and analyze such binding behavior easily. On comparing the overlap events detected by **bedtools intersect** and **bindcompare**, Bedtools returns 40.64% of the events detected by BindCompare (**Figure S4**). In the inverse direction, **bindcompare** returns 146.08% additional events in comparison to those detected by **bedtools intersect**. The set returned by **bedtools intersect** is a subset of those returned by **bindcompare**.

In contrast to BindCompare, DiffBind is better suited for testing hypotheses related to specific conditions or sets of proteins. In particular, neither DiffBind nor Bedtools easily integrates and compares large numbers of protein-nucleic acid data sets to generate new hypotheses. Visualizations produced by bindexplore streamline this process and can identify potential regulatory networks. These comprehensive and reproducible overlap analyses, powered by unique visualization, differentiate BindCompare from DiffBind and Bedtools.

Furthermore, BindCompare has a unique focus on user accessibility. Its GUI interface allows users to run analyses without a background in scripting, which is necessary for bedtools and DiffBind. With the rise of new computing platforms, like Apple Silicon, many R/Bioconductor packages cannot be run without docker images or special compute environments. This barrier can make running exploratory analysis to generate new hypotheses challenging. BindCompare is compatible with any architecture (x86_64, Apple Silicon, etc.) through its PyPI (pip) installation.

Overall, BindCompare is a unique tool that allows rigorous, reproducible binding analysis, enabling users to develop hypotheses for downstream analysis. BindCompare occupies an essential niche in bioinformatics by providing a user-friendly approach to generating new hypotheses, for which tools such as DiffBind and bedtools have not been explicitly designed.

#### 1.8: Data Availability

BindCompare is an open-source package that is available on the Python Packaging Index (PyPI, https://pypi.org/project/bindcompare/) with the source code available on GitHub (https://github.com/pranavmahabs/bindcompare). Complete documentation for the package can be found at both of these links.

This study leveraged existing ChIP-seq and eCLIP datasets from *the H. Sapiens* and *D. melanogaster* projects (**Supplementary File 3, Table 2**). All human ChIP-seq and eCLIP datasets, aligned to the hg19 genome build, can be found on GEO with accession number GSE77634. The ChIP-seq experiments for Drosophila datasets can also be found on GEO with accession numbers GSE133637, GSE220053, GSE205987, and GSE174781.
