## Supplemental File 2: Figures for "BindCompare: A Novel Integrated Protein-Nucleic Acid Binding Analysis Platform"

### Supplementary File 2

#### Figure S1

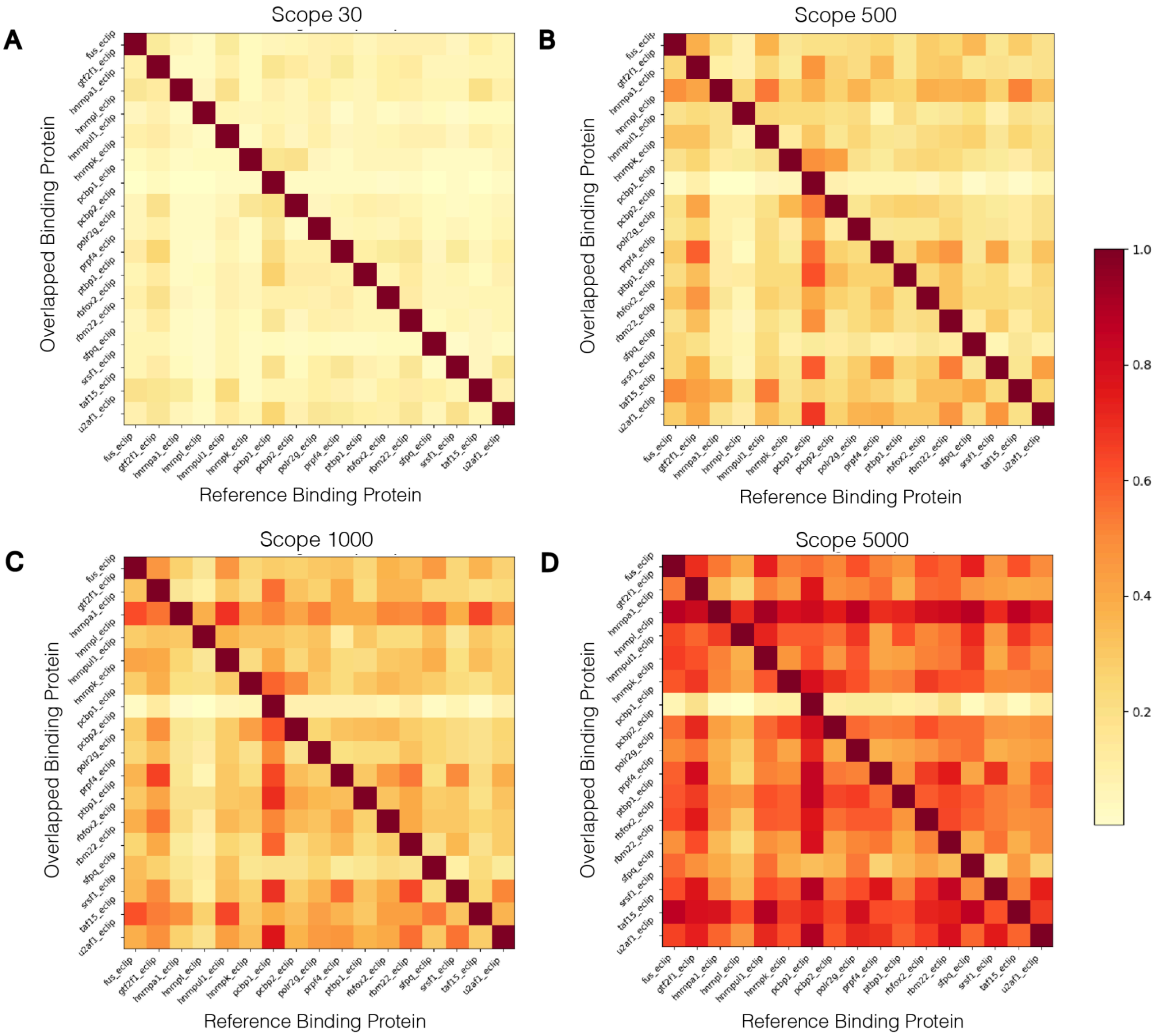

#### Figure S2

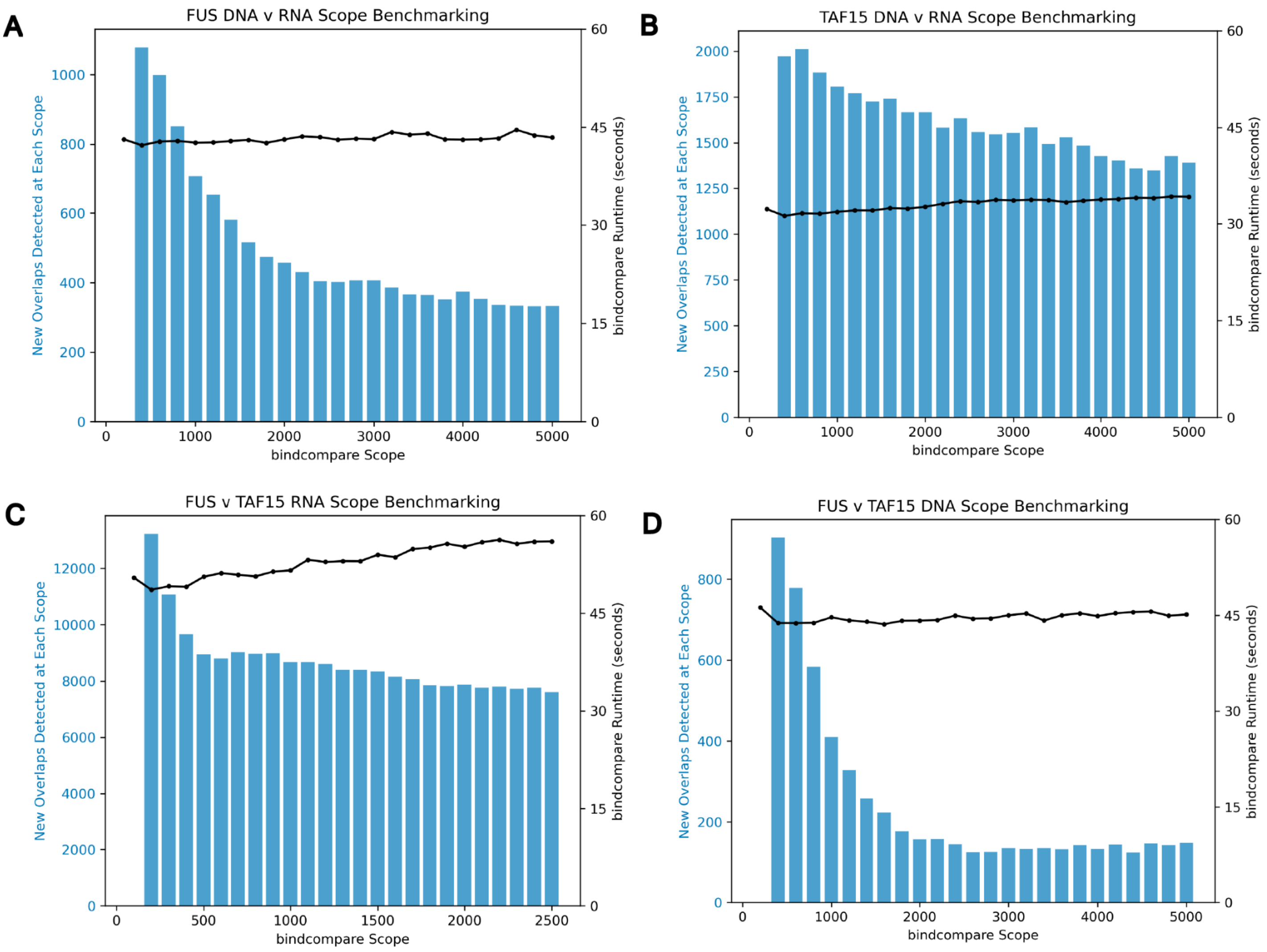


#### Figure S3

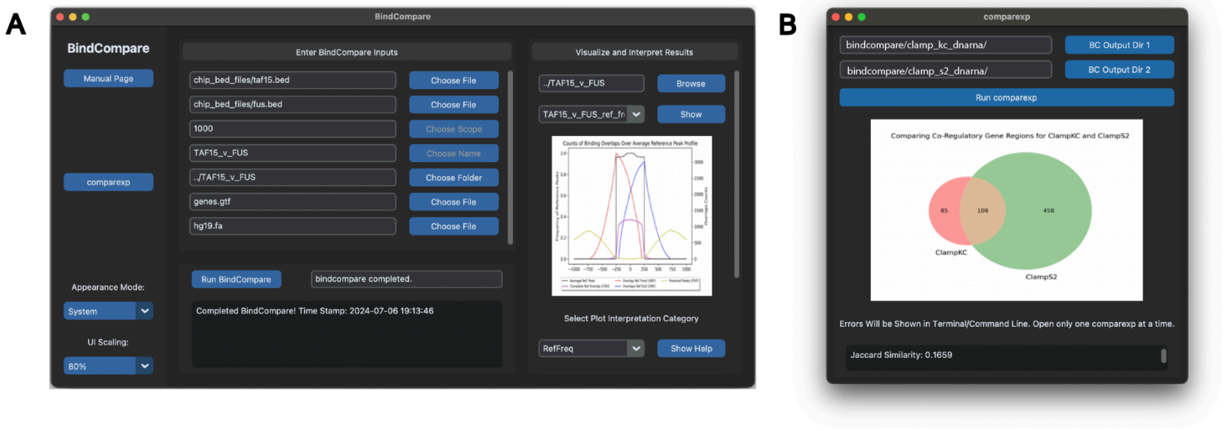

#### Figure S4

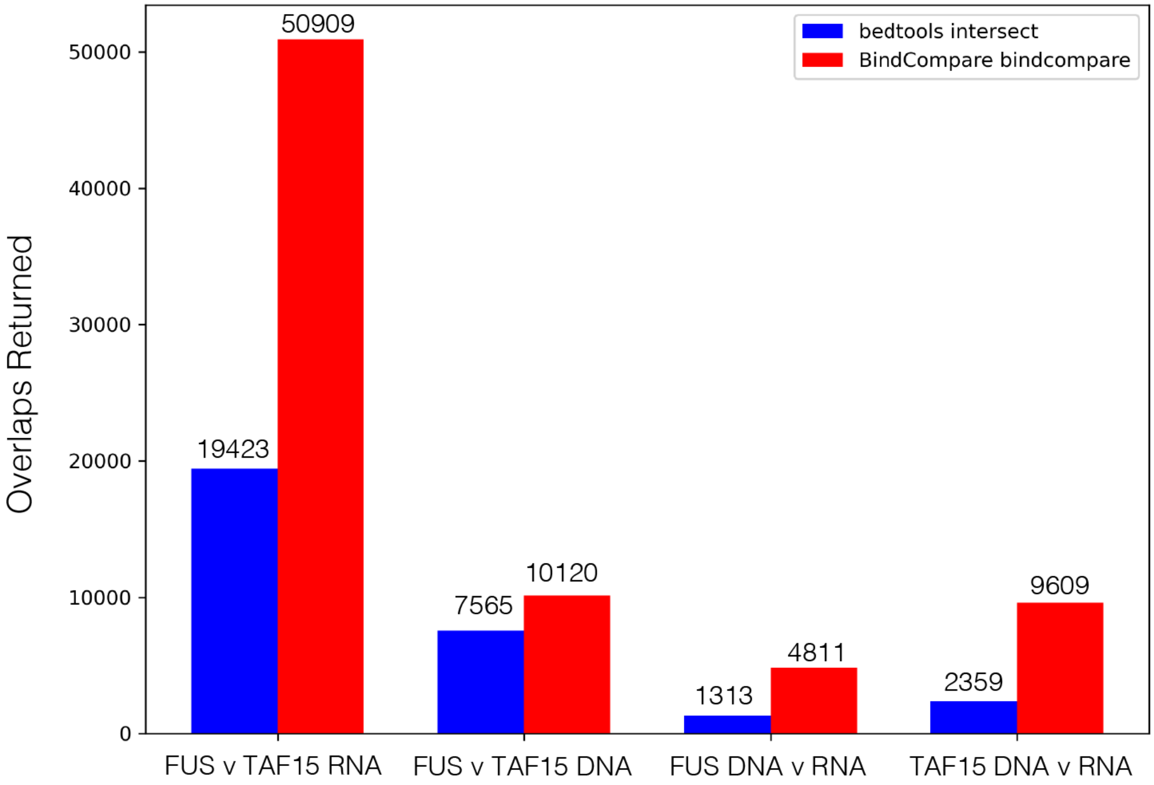

bedtools intersect
BindCompare bindcompare
Overlaps Returned
0
10000
20000
30000
40000
50000
19423
50909
7565
10120
1313
4811
2359
9609
FUS v TAF15 RNA
FUS v TAF15 DNA
FUS DNA v RNA
TAF15 DNA v RNA

#### Figure S5

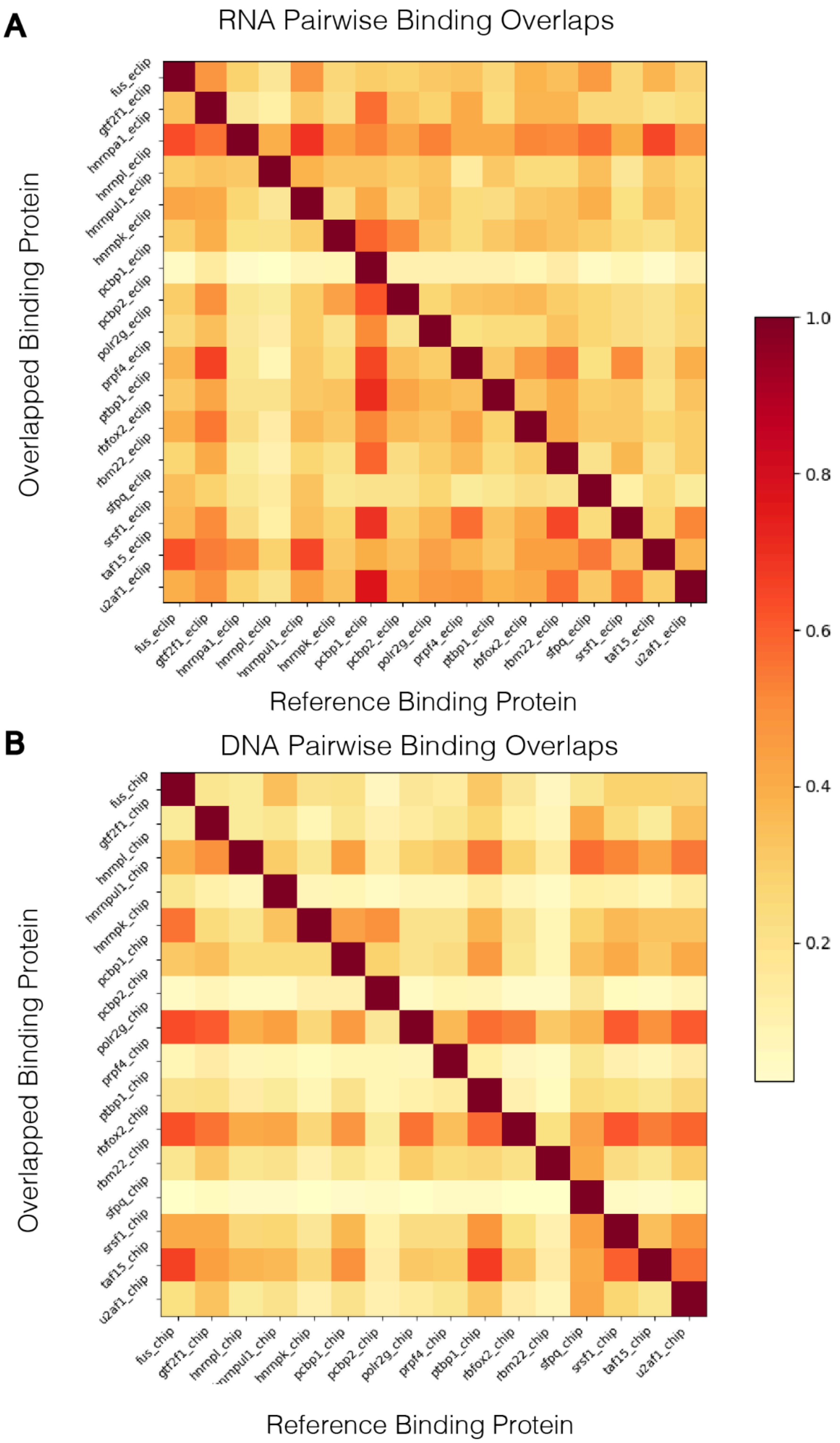
A
RNA Pairwise Binding Overlaps
Overlapped Binding Protein
Reference Binding Protein
fus_eclip
gtf2f1_eclip
hnrnpa1_eclip
hnrnpl_eclip
hnrnpul1_eclip
hnrnpk_eclip
pcbp1_eclip
pcbp2_eclip
polr2g_eclip
prpf4_eclip
ptbp1_eclip
rbfox2_eclip
rbm22_eclip
sfpq_eclip
srsf1_eclip
taf15_eclip
u2af1_eclip
B
DNA Pairwise Binding Overlaps
Overlapped Binding Protein
Reference Binding Protein
fus_chip
gtf2f1_chip
hnrnpl_chip
hnrnpul1_chip
hnrnpk_chip
pcbp1_chip
pcbp2_chip
polr2g_chip
prpf4_chip
ptbp1_chip
rbfox2_chip
rbm22_chip
sfpq_chip
srsf1_chip
taf15_chip
u2af1_chip
1.0
0.8
0.6
0.4
0.2

### Figure S6

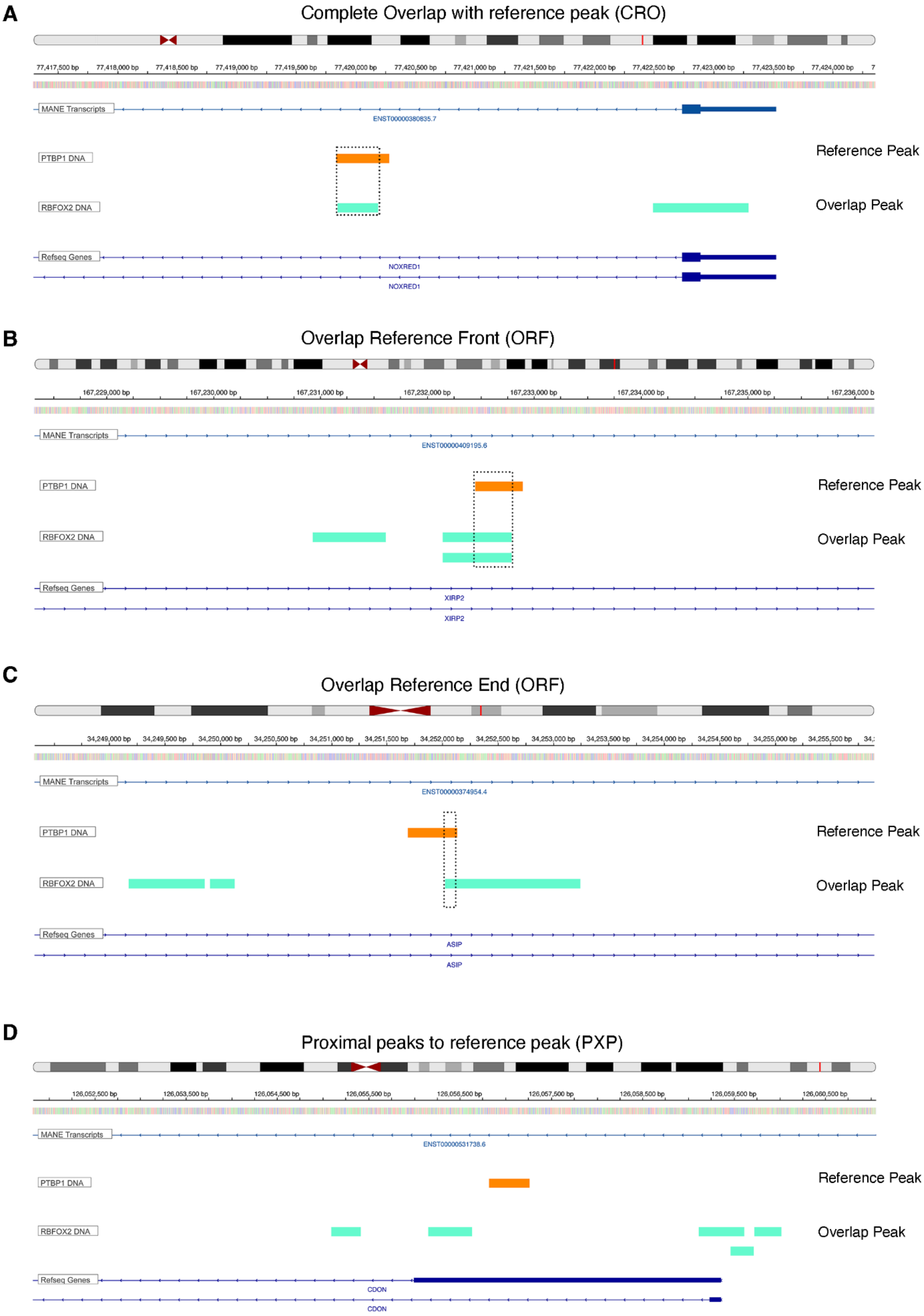
A
Complete Overlap with reference peak (CRO)
MANE Transcripts
ENST00000380835.7
PTBP1 DNA
Reference Peak
RBFOX2 DNA
Overlap Peak
Refseq Genes
NOXRED1
NOXRED1
B
Overlap Reference Front (ORF)
MANE Transcripts
ENST00000409195.6
PTBP1 DNA
Reference Peak
RBFOX2 DNA
Overlap Peak
Refseq Genes
XIRP2
XIRP2
C
Overlap Reference End (ORF)
MANE Transcripts
ENST00000374954.4
PTBP1 DNA
Reference Peak
RBFOX2 DNA
Overlap Peak
Refseq Genes
ASIP
ASIP
D
Proximal peaks to reference peak (PXP)
MANE Transcripts
ENST00000531738.6
PTBP1 DNA
Reference Peak
RBFOX2 DNA
Overlap Peak
Refseq Genes
CDON
CDON

Figure S7

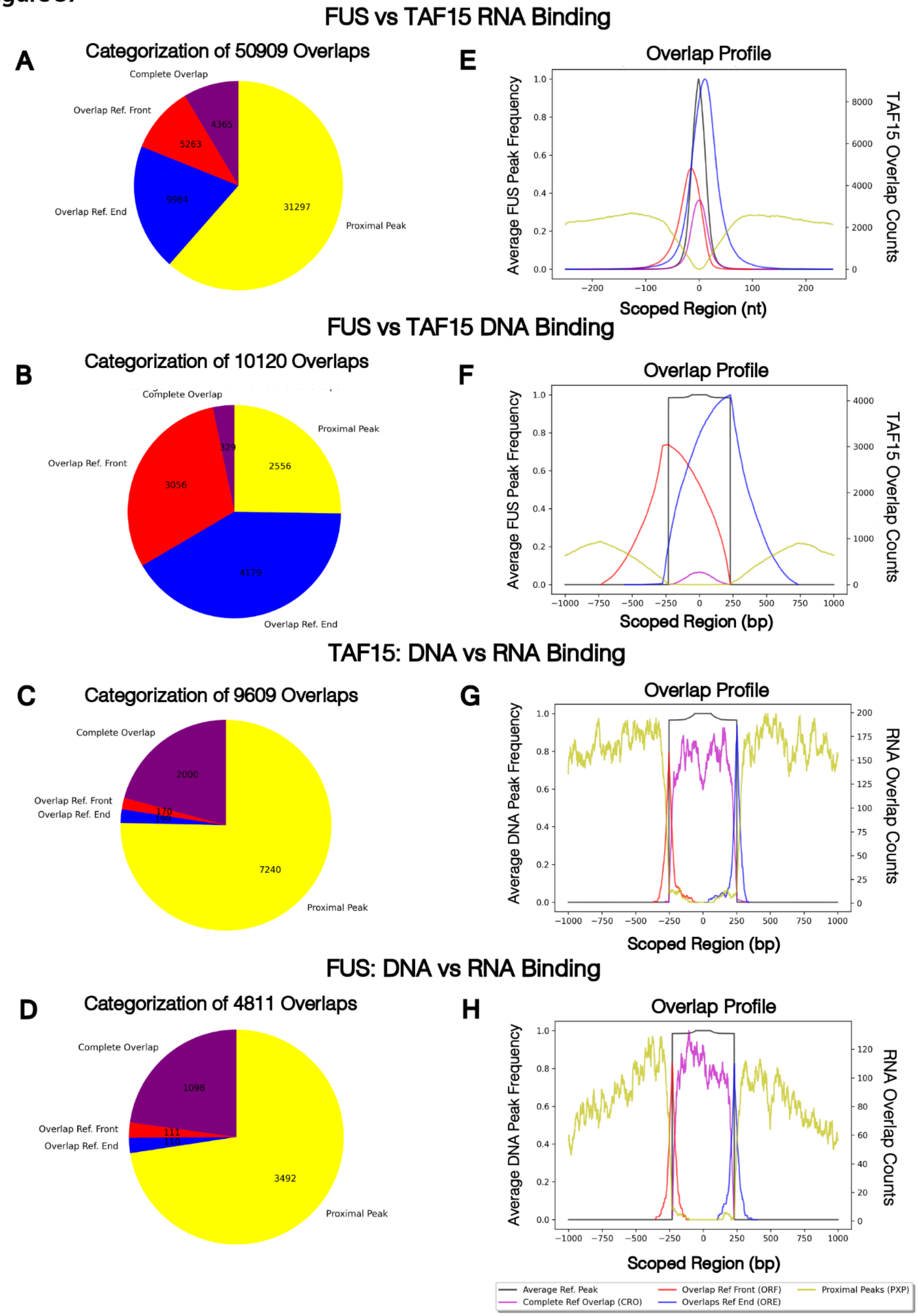
FUS vs TAF15 RNA Binding
A
Categorization of 50909 Overlaps
Complete Overlap
Overlap Ref. Front
Overlap Ref. End
Proximal Peak
4365
5263
9984
31297
E
Overlap Profile
Average FUS Peak Frequency
TAF15 Overlap Counts
Scoped Region (nt)
FUS vs TAF15 DNA Binding
B
Categorization of 10120 Overlaps
Complete Overlap
Proximal Peak
Overlap Ref. Front
Overlap Ref. End
329
2556
3056
4179
F
Overlap Profile
Average FUS Peak Frequency
TAF15 Overlap Counts
Scoped Region (bp)
TAF15: DNA vs RNA Binding
C
Categorization of 9609 Overlaps
Complete Overlap
Overlap Ref. Front
Overlap Ref. End
Proximal Peak
2000
170
7240
G
Overlap Profile
Average DNA Peak Frequency
RNA Overlap Counts
Scoped Region (bp)
FUS: DNA vs RNA Binding
D
Categorization of 4811 Overlaps
Complete Overlap
Overlap Ref. Front
Overlap Ref. End
Proximal Peak
1098
111
110
3492
H
Overlap Profile
Average DNA Peak Frequency
RNA Overlap Counts
Scoped Region (bp)
Average Ref. Peak
Complete Ref Overlap (CRO)
Overlap Ref Front (ORF)
Overlaps Ref End (ORE)
Proximal Peaks (PXP)

#### Figure S8

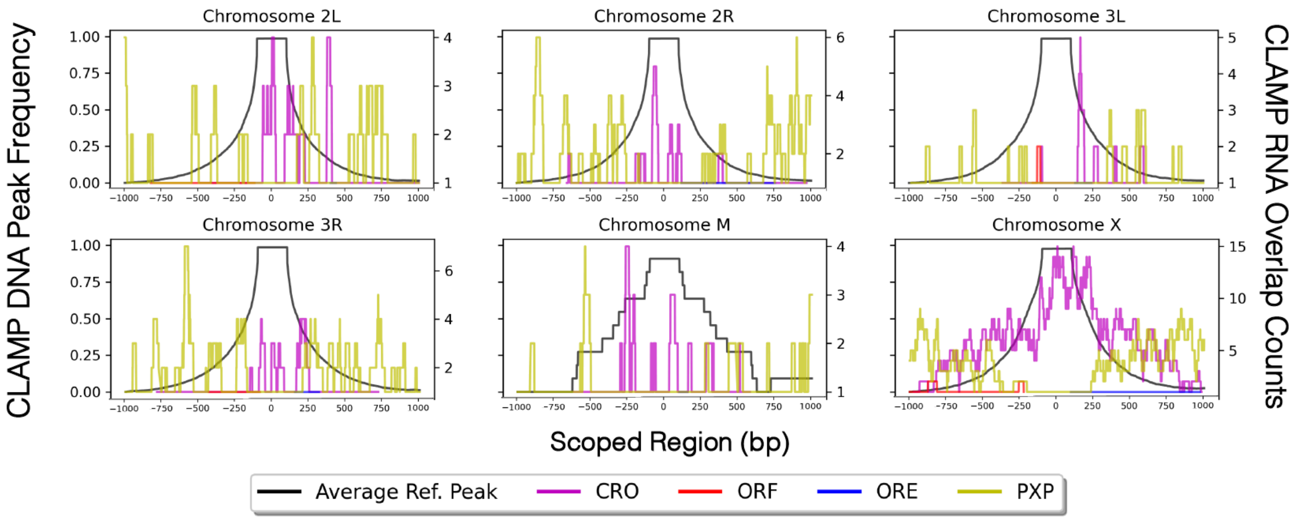


#### Figure S9

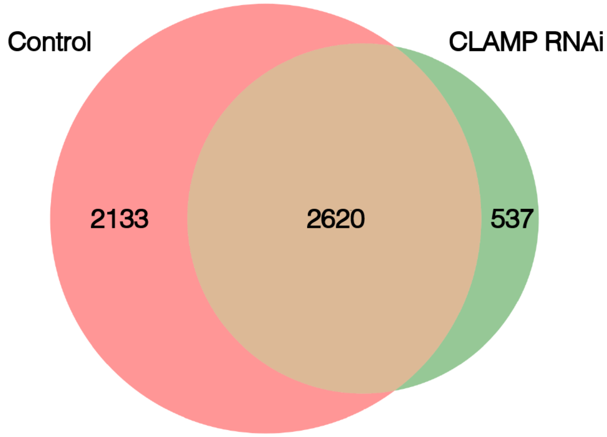

Figure S1 **Impact of scope on bindexplore results. bindexplore** generated a heat-map matrix comparing eCLIP data for 16 human RBPs, pair-wise comparison of genome-wide common RNA targets between each pair with varying scopes/bin sizes: **A** 30 bp, which is also the average peak size of the eCLIP peaks, **B** 500 bp, **C** 1000 bp (the default bin size), and **D** 5000 bp bin sizes. These four runs had respective runtimes of 3.345, 2.996, 2.970, and 2.845 seconds. Deep Red = correlation value of 1 (Complete overlap), orange shades = correlation value between 1 and 0.5, and yellow shades = correlation value between 0.5 and 0.

Figure S2 **Impact of scope on bind compare's identified overlaps and runtime. A-D** Bar plots show along the right-hand side y-axis the number of new overlapping events detected at different scope ranges mentioned along the x-axis and the time taken to run each set is shown in the left-handside y-axis. FUS ChIP-seq peaks compared to FUS eCLIP (**A**), TAF15 ChIP-seq peaks compared to TAF15 eCLIP (**B**), FUS eCLIP peaks compared to TAF15 eCLIP (**C**), and FUS ChIP-seq peaks compared to TAF15 ChIP-seq (**D**). Comparisons involving DNA overlaps (**A, B,** and **D**) were tested on scopes ranging from 200-5000bp (inclusive, incremented by 200). In contrast, comparison for RNA-RNA overlaps (**C**) was tested on scopes ranging from 100-2600 (inclusive, incremented by 100).

Figure S3 **BindCompare GUI interface. A.** The **bindcompare** app window launches from the command line. Here, **bindcompare** can be run and interpreted using the provided help text. **B**. The **comparexp** interface launches from the main BindCompare app. This interface allows you to run all the aforementioned commands except for **bindexplore**. When running scripts with the GUI, any messages produced in the terminal will also be piped into the GUI interface, informing users of potential file errors and indicating when the script is in progress or completed.

Figure S4 **Comparing bindcompare and bedtools intersect.** Bar plots showing the number of overlapping events detected from **bedtools intersect** (**blue bar**) with the number of overlapping events returned from **bindcompare** for four sets of comparisons along the x-axis. **The numbers** at the top of each bar denote the overlaps detected by each method.

Figure S5 **bindexplore outputs comparing human RNA-binding proteins binding to nucleic acids. A bindexplore** generated a heat-map matrix comparing eCLIP data for 16 human RBPs with a bin size of 1000, pair-wise comparison of genome-wide common RNA targets between each pair. **B bindexplore** generated a heat-map matrix comparing ChIP data for 16 human RBPs with a bin size of 1000, pair-wise comparison of genome-wide common DNA targets between each pair. Red = correlation value of 1 (Complete overlap), orange shades = correlation value between 1 and 0.5, and yellow shades = correlation value between 0.5 and 0.

Figure S6 **Peak overlap categories identified in bindcompare**. **A-D** IGV browser screenshots show different categories of overlap between protein-nucleic acid reference peaks and those of protein-nucleic acid peaks compared using the **bindcompare** module. The dotted rectangular box denotes the region of overlap.

Figure S7 **bindcompare outputs comparing Fus and Taf15 nucleic acid binding**. **A-D** Pie-charts show the number of peak overlaps between reference and test nucleic acid binding proteins in each category: purple = complete peak overlaps (CRO), red = 5'overlap at the front of the reference peak (ORF), blue = 3'overlap at the end of the reference peak (ORE), yellow = peaks proximal to the reference peak (PXP) ($\pm$1000bp). **E-H** Frequency plots (overlap profile) show the counts of each overlap event over the scoped domain (x-axis). A purple line for complete peak overlaps (CRO), red for 5'overlap at the front of the reference peak (ORF), blue for 3'overlap at the end of the reference peak (ORE), and yellow for peaks proximal to the reference peak. The average reference peak frequency at the scoped region is shown in black. The dip in the yellow line (**E-H**) represents the proximal peaks, i.e., those that do not completely or partially overlap but are present within the scope region of $\pm$ 1000bp (scope = 1000bp). In **G** and **H**, the complete overlap RNA peaks (purple line) are concentrated at the middle of the DNA peak distribution (black line), indicating that the Fus/Taf15 protein binds to both DNA and RNA at similar positions. The partial RNA peak overlaps at the front (red line) and end (Blue lines), as expected, and is peaking at the front and the end of the DNA peak distribution (black line).
In contrast, the non-overlapping RNA proximal peaks (yellow line) show a dip at the center of the DNA peak distribution and are present outside it, mainly flanking it. The completely overlapping peak dips in the middle of **F** because we are comparing two different DNA-binding proteins, and the likelihood of two DNA-binding proteins binding in the exact genomic location is lower. Since TFs interacting in the same area might bind at adjacent sites on the DNA, so we see a higher frequency for red and blue partially overlapping peaks at the front and end of the reference peak in **F**.

Figure S8 **Distribution of CLAMP TF binding with DNA and RNA across individual chromosomes.** The aggregate overlap profile visualizes the frequency of overlapping events over the scoped domain across individual chromosomes, with most overlap events occurring on the X chromosome.

Figure S9 **Clamp regulates MLE-DNA targets.** Venn diagram output from **comparexp** compares common genomic loci associated with CLAMP and MLE-DNA targets in the presence (red circle) and absence (green circle) of the CLAMP transcription factor. New MLE-DNA targets (absence of CLAMP) are in green (N=537), and retained MLE-DNA targets are in brown (N=2620). N denotes the number of genes associated with MLE-DNA peaks. The gene lists had a Jaccard similarity coefficient score of 0.4953, indicating less similarity between the groups.
